# Asymmetric redundancy of *ZERZAUST* and *ZERZAUST HOMOLOG* in different accessions of *Arabidopsis thaliana*

**DOI:** 10.1101/452615

**Authors:** Prasad Vaddepalli, Lynette Fulton, Kay Schneitz

## Abstract

Divergence among duplicate genes is one of the important sources of evolutionary innovation. But, the contribution of duplicate divergence to variation in Arabidopsis accessions is sparsely known. Recently, we studied the role of a cell wall localized protein, ZERZAUST (ZET), in Landsberg *erecta* (L*er*) accession. Here, we present the study of *ZET* in Columbia (Col) accession, which not only showed differential expression patterns in comparison to L*er*, but also revealed its close homolog, *ZERZAUST HOMOLOG (ZETH)*. Although, genetic analysis implied redundancy, expression analysis revealed divergence, with *ZETH* showing minimal expression in both Col and L*er*. In addition, *ZETH* shows relatively higher expression levels in Col compared to L*er*. Our data also reveal compensatory up-regulation of *ZETH* in Col, but not in L*er*, implying it is perhaps dispensable in L*er*. However, a novel CRISPR/Cas9-induced *zeth* allele confirmed that *ZETH* has residual activity in L*er*. The results provide genetic evidence for accession-specific differences in compensation mechanism and asymmetric gene contribution. Thus, our work reveals a novel example for how weakly expressed homologs contribute to diversity among accessions.

## Introduction

How genetic variation translates into phenotypic variation is of immense scientific interest (Weigel 2012). Among others, gene duplication followed by functional divergence is an important source of evolutionary complexity and innovation in multicellular organisms (Ohno 1970; Lynch and Conery 2000; Lynch and Katju 2004). Many gene pairs after duplication revert to single gene state but the ones that sustained undergo functionalization. The Arabidopsis genome underwent several duplication events which resulted in large number of homologous genes and regions across the genome (Blanc and Wolfe 2004; Ambrosino *et al*. 2016; Panchy *et al*. 2016). The functional importance of homologs has been demonstrated in various aspects of plant signaling and metabolism (Briggs *et al*. 2006). But, whether and how the differentiation in duplicate gene expression contributes to accession variation in Arabidopsis is not known.

Studies have shown that divergence of many duplicate genes occurs by expression divergence among and within species (Gu *et al*. 2004; Li *et al*. 2005). This phenomenon expands gene regulatory networks and contributes to physiological and morphological diversity (Carroll 2000; Lynch and Conery 2000; Gu *et al*. 2004; Rensing 2014). In Arabidopsis, about two-thirds of duplicates were shown to exhibit expression divergence (Haberer *et al*. 2004). An evolutionary study on gene duplication revealed that duplicate genes show a high degree of variance in expression within species and suggested that this variation partly depends upon the biological function of the gene involved (Kliebenstein 2008). Another study also found high variance of duplicated gene expression between closely related *A. thaliana* and *A. arenosa* (Ha *et al*. 2009).

Functional redundancy among homologs is widespread in Arabidopsis, since several single loss-of-function mutants lack phenotype (Briggs *et al*. 2006). Homologous genes can be either fully or partially redundant. But, when two homologous genes show unequal genetic redundancy, a mutation in one of them causes a phenotype and the phenotype is enhanced when the other homolog is mutated as well. Interestingly, the defect in the other homolog itself doesn't result in any phenotype on its own. For example, the receptor-like kinase gene *BRASSINOSTEROID INSENSITIVE 1 (BRI1)* is accompanied by its close homolog *BRI1-LIKE1 (BRL1)*. Although *brl1* lacks a mutant phenotype it enhances the severe dwarf phenotype of *bri1* mutants (Caño-Delgado *et al*. 2004). This kind of unequal functional redundancy can be explained by divergence in duplicate expression, however, their perseverance in plant genome is under debate given the dispensable nature of the duplicate.

Genetic factors involved in plant morphogenesis will have crucial role in the differentiation of various Arabidopsis accessions. Tissue morphogenesis in Arabidopsis requires the cell wall-localized GPI-anchored β-1,3 glucanase ZERZAUST (ZET) (Vaddepalli *et al*. 2017). *ZET* was initially identified as a genetic component of the *STRUBBELIG (SUB)* signaling pathway along with QUIRKY, a C2 domain containing protein (Fulton *et al*. 2009). Absence of *ZET* results in a so-called *strubbelig-like mutant* (*slm*) phenotype characterized by abnormal integument initiation and outgrowth, aberrant floral organ and stem morphology, reduced plant height and irregular leaf shape.

Our previous studies have shown that mutations in *SUB* and *QKY* in Col background result in obvious *slm* mutant phenotypes (Fulton *et al*. 2009; Vaddepalli *et al*. 2011, 2014). But in the current work, we discovered that *ZET* acts differently in Col accession due to the presence of the close homolog *ZERZAUST HOMOLOG* (*ZETH*). Using genetic and gene expression tools we show how *ZET* and *ZETH* diverged between the two common laboratory accessions Col and *Ler* in terms of expression and function. Furthermore, we try to understand the contribution of the weakly expressing redundant homolog on the morphological diversity of accessions.

## Results

### Molecular identification of *ZETH*

In *Ler* background, *zet-1* carrying a loss-of-function mutation in *ZET* locus (At1g64760) (Fig. 1A,B), shows a strong *slm* mutant phenotype (Vaddepalli *et al*. 2017) (Fig. 1D,F). Except one amino acid in the signal peptide, ZET shows no difference between Col and *Ler*. We investigated the functionality of *ZET in* Columbia accession by analyzing two available T-DNA insertion lines (*zet-3* and *zet-4*) (Fig. 1A). We expected the T-DNA insertion in *zet-4* to cause a mutant phenotype as it is predicted to result in a truncated ZET protein (Fig. 1A,B). But, the plants surprisingly failed to display the twisted morphology, characteristic of *slm* mutants (Fig. S1B). We asked, if the observed accession-specific phenotypic differences could relate to a close homolog of *ZET*. A BLAST search with the *ZET* coding sequence revealed that *ZET is* most closely related to At2g19440 with 89 percent identity at the amino acid level (Fig. S1A). We named this gene *ZERZAUST HOMOLOG (ZETH). ZET* and *ZETH* form a subclade within the larger β clade of β-1,3 glucanase (BG) genes, which comprises 11 members (Doxey *et al*. 2007; Gaudioso-Pedraza and Benitez-Alfonso 2014). Sequence-based analysis of the evolutionary history revealed that ZET and ZETH duplication is specific to species within the Arabidopsis lineage (Fig. S2).

**Fig. 1.**
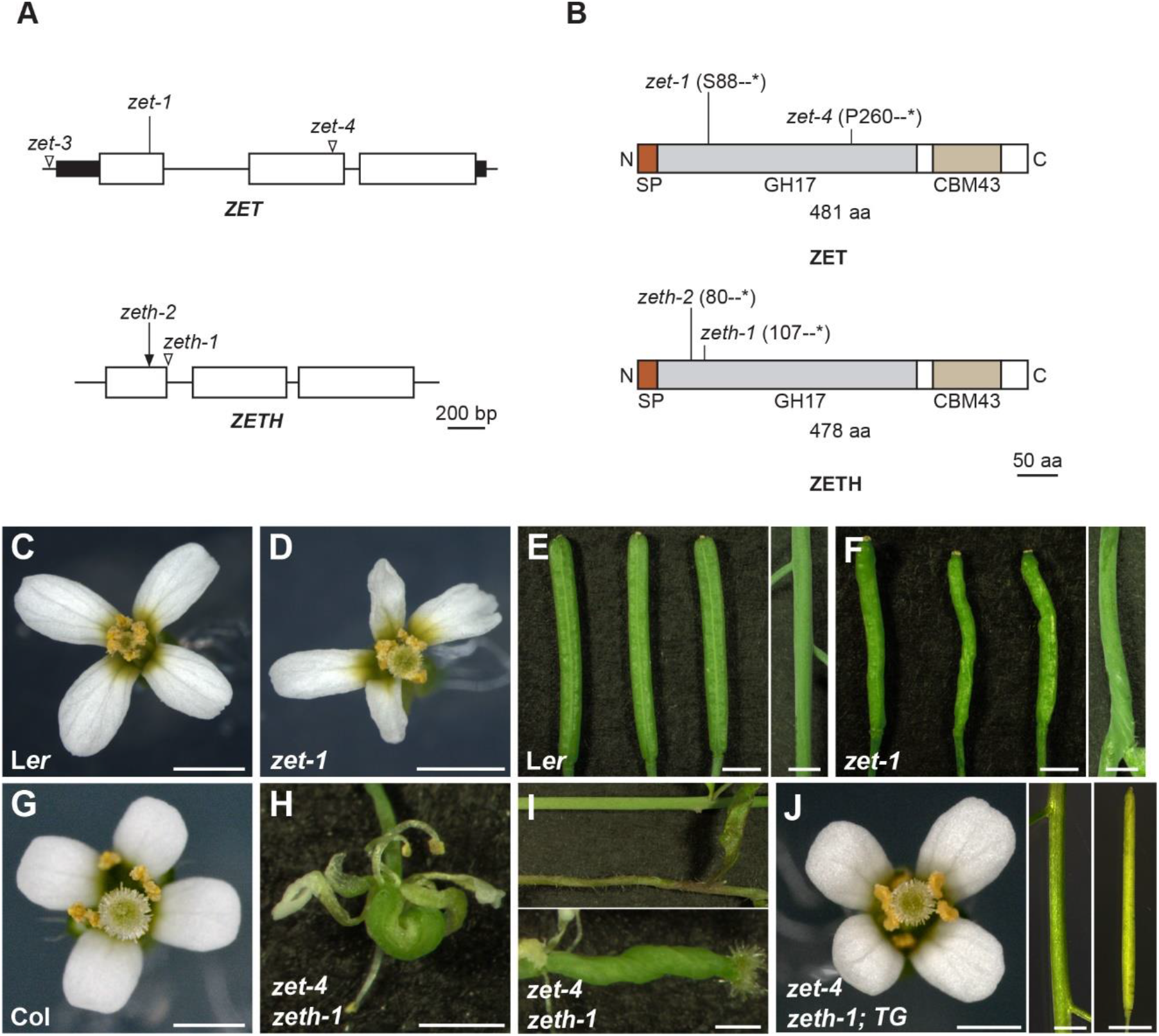
Molecular and genetic characterization of *ZET* and *ZETH*. (A) Cartoon depicting the genomic organization of *ZET* and *ZETH*. Horizontal lines represent introns. Filled rectangles mark untranslated regions. The positions of EMS-, CRISPR-, and T-DNA-induced mutations are denoted by lines, arrow, and open triangles, respectively. (B) Schematic view of the predicted ZET protein. The signal peptide (SP), GH17 and CBM43/X8 domains are highlighted. The position and effects of the *zet* mutations are indicated. Phenotypic analysis (C-J). Comparison of (C, E) L*er*. (D, F) *zet-1*, (G) Col and (H, I) *zet-4 zeth-1*. Note the aberrant flower (D, H), stem and silique (F, I) morphology of mutants. Mutant phenotype is strong in *zet-4 zeth-1*double mutant (H, I). (J) *zet-4 zeth-1* double mutant phenotype is complemented by *pZET::TS:ZET* (*TG*). Scale bars:1 mm

To assess ZETH activity in Col, we investigated a T-DNA line (*zeth-1*), which is presumed to carry a truncated protein (Fig. 1A,B). But, like *zet-4*, the *zeth-1* insertion line also failed to show a mutant phenotype (Fig. S1C). However, the *zet-4* and *zeth-1* double mutant exhibited a strong phenotype (Fig. 1G-I, S1D) suggesting that the two genes act redundantly. The double mutant resembled *zet-1* plants except for appearing less bushy but with exaggerated twisting of flowers and increased sterility. Nevertheless, we could complement the double mutant plants by introducing a construct encoding a translational fusion of ZET to the GFP variant T-Sapphire driven by the endogenous *ZET* promoter (pZET::TS:ZET), ruling out the contribution of any other background mutation (Fig. 1J). This construct was used in a previous study to complement *zet-1* mutants in L*er* background (Vaddepalli *et al*. 2017).

### Accession-specific regulation of *ZET* and *ZETH* expression

Next, we analyzed the expression pattern of *ZET* and *ZETH* in Col and L*er* accessions to assess the cause for the mutant phenotype disparities between the two accessions. Our qPCR data from various tissues revealed that these genes are co-expressed (Fig. 2A). Surprisingly, we found much lower levels of *ZETH* transcripts in comparison to *ZET*. Moreover, *ZETH* expression was even further reduced in L*er* where it was barely detectable in rosette leaves, stems, or flowers. Additional qPCR tests revealed that *ZET* and *ZETH* expression levels in seedlings and flowers undergo compensatory regulation in the *zeth-1* and *zet-4* mutants in Col, respectively (Fig. 2B). This result provides evidence for redundant functions of *ZET* and *ZETH* in the Col background and thus offers a convenient explanation for the lack of phenotype in single mutants in this accession. Interestingly, this compensatory regulation appears to be absent in flowers of L*er* accession since *ZETH* expression was not detectably upregulated in *zet-1* mutant.

**Fig. 2.**
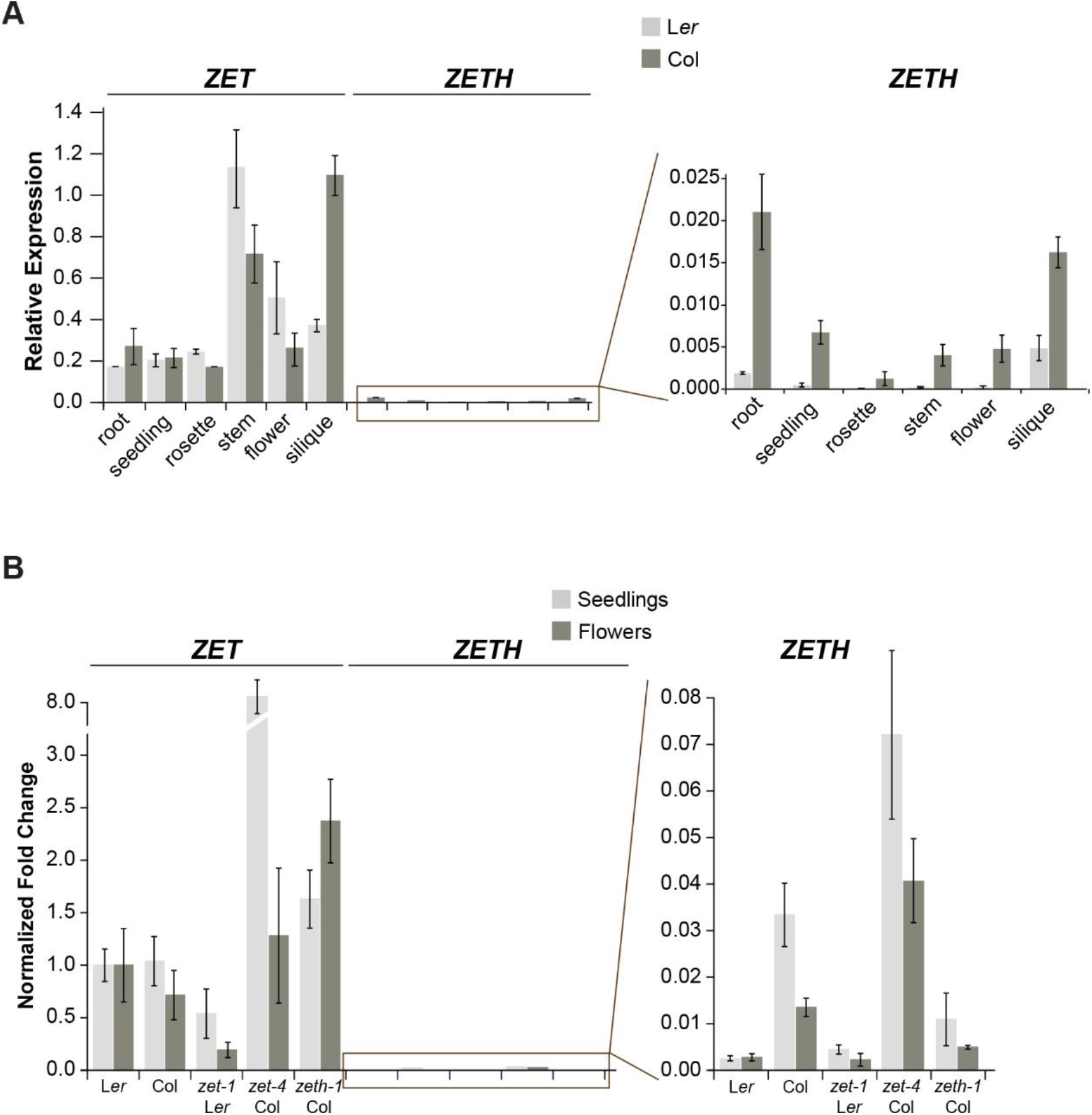
Expression analysis of *ZET* and *ZETH*. (A) Tissue distribution of *ZET* and *ZETH* transcript expression levels by qPCR. (n= 3 biological replicates). Means ± SEMs are indicated. Age of plants: roots and seedlings, 10 days; rosette, 3 weeks; stem, flowers (stages 1-12) and siliques (stage 17), 5 weeks. (B) Comparison of *ZET* and *ZETH* mRNA levels in seedlings and stage 1-12 flowers of indicated mutants by qPCR. (n= 3 biological replicates). Means ± SEMs are indicated.

### L*er* carries a functional *ZETH* gene

Despite minimal *ZETH* expression profiles in both accessions, appearance of prominent phenotypes only in L*er*, when *zet* is mutated, can be attributed to any or all of the following reasons. It could be because of the low expression of *ZETH*, the lack of compensatory mechanism, or the L*er* version of ZETH exhibiting a different amino acid composition when compared to Col which might affect its activity. The L*er*/Col variants of ZET differ by only one amino acid at position 3 in the predicted signal peptide (change from an asparagine to a lysine) but there are nine amino acid differences between the L*er*/Col variants of ZETH (Fig 3A). We wanted to test if these changes affect the activity of ZETH in L*er*. For this purpose, we replaced the *ZET* coding sequence 3’ to the predicted signal peptide with the equivalent L*er* or Col variants of *ZETH* sequence in our complementing pZET::TS:ZET reporter (Vaddepalli *et al*. 2017). This resulted in *zet-1* plants transgenic for the L*er* or Col variants of *ZETH*, under the control of the native *ZET* promoter (*pZET::TS:ZETHL/C zet-1*). Interestingly, the T1 plants of the transgenic *zet-1* lines exhibited a wild-type phenotype with both variants (Figures 3B-G) (80/80 (*TS:ZETHL*), 167/172 (*TS:ZETHC*)) indicating that the accession-specific amino acid alterations do not affect ZETH function.

**Fig. 3.**
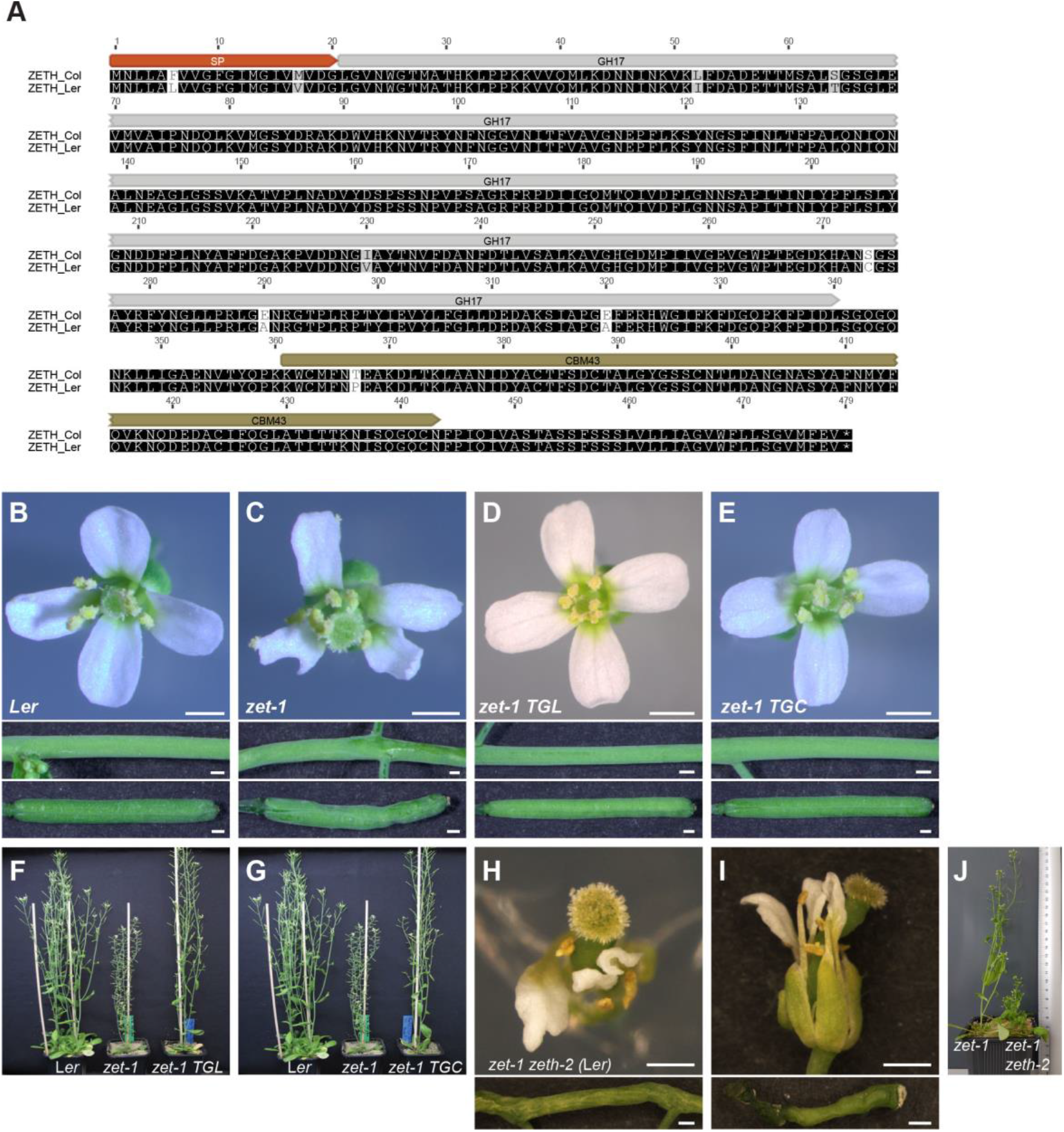
*ZETH* in L*er* has residual function. (C) Alignment of ZETH amino acid sequences of Col and L*er*. (B-E) Complementation of *zet-1* phenotype by two TS:ZET reporter constructs. Upper panels: Stage 13 flower. Middle panel: siliques. Bottom panel: Stem. Genotypes are indicated. (C) Note aberrant floral morphology. Siliques and stems are twisted. (D, E) Note normal phenotype of a *zet-1* plant carrying either the L*er* or Col variant of *ZETH* under the control of the endogenous *ZET* (L*er*) promoter (TGL: *pZET::TS:ZETH*/L*er*, TGC: *pZET::TS:ZETH*/Col). (F, G) Whole-plant appearance. Genotypes are indicated. (H-J) Phenotype of *zet-1 zeth-2* (L*er*) mutant. Mutant phenotype is exaggerated. Scale bars: 0.5 mm.

Although *ZETH* of L*er* is functional, its expression is quite weak indicating its functional contribution is perhaps insignificant. But, *zet1 zeth-1* double mutants in Col accession exhibit a stronger phenotype compared to *zet-1* (L*er*) (Fig. 1G-J). These interesting observations prompted us to check whether *ZETH* has some residual activity in L*er* even though its expression is very low. Using CRISPR\Cas9 technique (Wang *et al*. 2015) we generated a mutation in the first exon of *ZETH* in the *zet-1* background. The novel allele *zeth-2* is predicted to result in a truncated ZETH protein consisting of only the first 80 amino acids (Fig. 1A,B). Surprisingly, *zet-1 zeth-2* double mutant plants in L*er* showed an exaggerated *zet-1* phenotype and appeared closer to *zet-4 zeth-1* double mutants in the Columbia background (Fig. 3H-J). The finding indicates that the weakly expressed *ZETH* in L*er* exhibits residual activity, which is insufficient to fully substitute for the lack of *ZET*.

### *ZETH* acts in a dose dependent manner

Thus far, our results have established the functional role for the weakly expressed *ZETH* in both Col and L*er* accessions, implying small amount of ZETH protein can have a noticeable impact. Next, we asked if this gene is acting in a dose-dependent manner. For this purpose, we checked the effect of *ZETH* gene copy number on *zet* mutant phenotype (Fig. 4). Interestingly in the L*er* background, *zet-1 zeth-2/+* displayed an intermediate phenotype between the single *zet-1* mutant and the double *zet-1 zeth-2* mutant. The leaf petioles are somewhat elongated in *zet-1* with narrow blades. This phenotype got slightly enhanced in *zet-1 zeth-2/+* background, whereas *zet-1 zeth-2* double mutants show further worsening of the phenotype. The phenomenon of dosage-dependent enhancement of mutant phenotype was also observed for floral organs and was particularly obvious for siliques depending on the *ZETH* copy number. Surprisingly, in Columbia background the *zet-4 zeth-1/+* mutant showed wild type morphology in all the organs tested except siliques which displayed shortening of length with no twisting. Despite these peculiarities between accessions, the observations indicate that mutation in *zeth* contributes to the overall exaggerated morphologies of double mutants in a dose dependent manner. Our results also imply that the extent of the *ZETH* effect on plant morphology is accession dependent.

**Fig 4.**
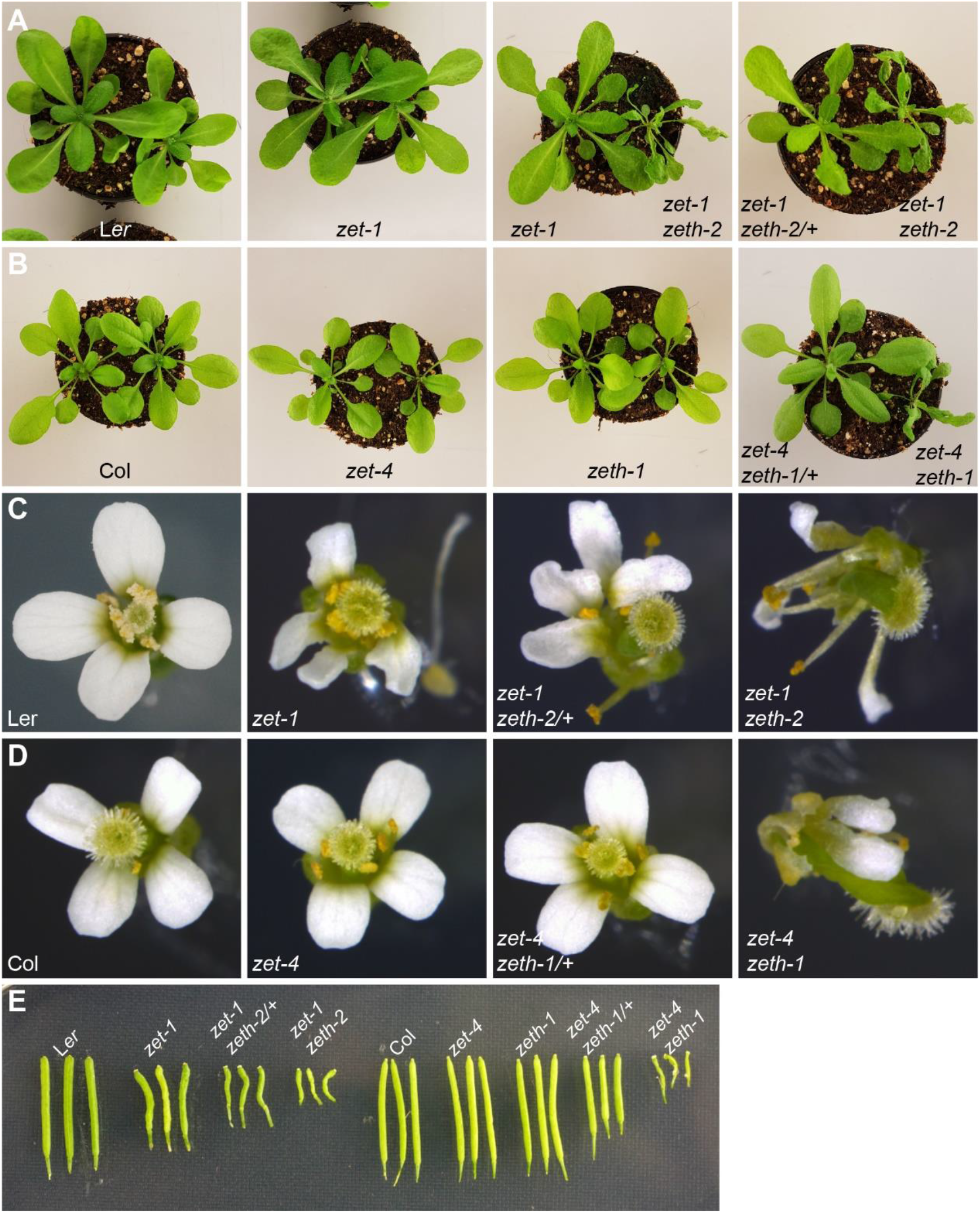
*ZETH* acts in dosage dependent manner. Genotypes are indicated. (A, B) Rosette leaves of three-week-old plants. (A) Notice slightly elongated petiole and narrow leaf blade phenotype in *zet-1*, which gets enhanced in *zet-1 zeth-2/+* and *zet-1 zeth-2* mutants. (B) In Col background aberrant morphology is apparent only in double mutants *zet-4 zeth-1*. (C, D) Flowers. (E) Siliques. Similar to leaves, twisting morphology of flowers and siliques is exaggerated progressively in L*er* mutants, but in Col only double mutant shows phenotype with the exception of siliques. Siliques of *zet-4 zeth-1/+* in Col are shorter.

### ERECTA influences the *zet-1* phenotype

Col and L*er* accessions display obvious discrepancies in their phenotypic appearance. For example, L*er* exhibits shorter stems and more compact inflorescences (Passardi *et al*. 2007). The phenotypic differences could be ascribed to genetic variability between the accessions and the contribution of accession-specific modifiers needs to be addressed. Major difference between the Col and L*er* accessions is the lack of *ERECTA* (*ER*) in the L*er* background. Previous work revealed that *SUB* and *QKY* show a synergistic interaction with *ERECTA* (*ER*) with respect to the control of plant height (Vaddepalli *et al*. 2011, 2014). We investigated the phenotype of *zet-1* (L*er*) plants transformed with pKUT196, a plasmid carrying 9.3 kb of Col-0 DNA spanning the entire genomic *ER* locus (Torii *et al*. 1996; Godiard *et al*. 2003). Like other *slms, zet-1 ER* transgenic plants also exhibited ameliorated plant height and stem twisting whereas aberrant floral organ phenotype appeared unaffected by the *ER* transgene (Fig. 5). Our result exemplifies the influence of accession-specific modifiers on plant morphology.

**Fig. 5.**
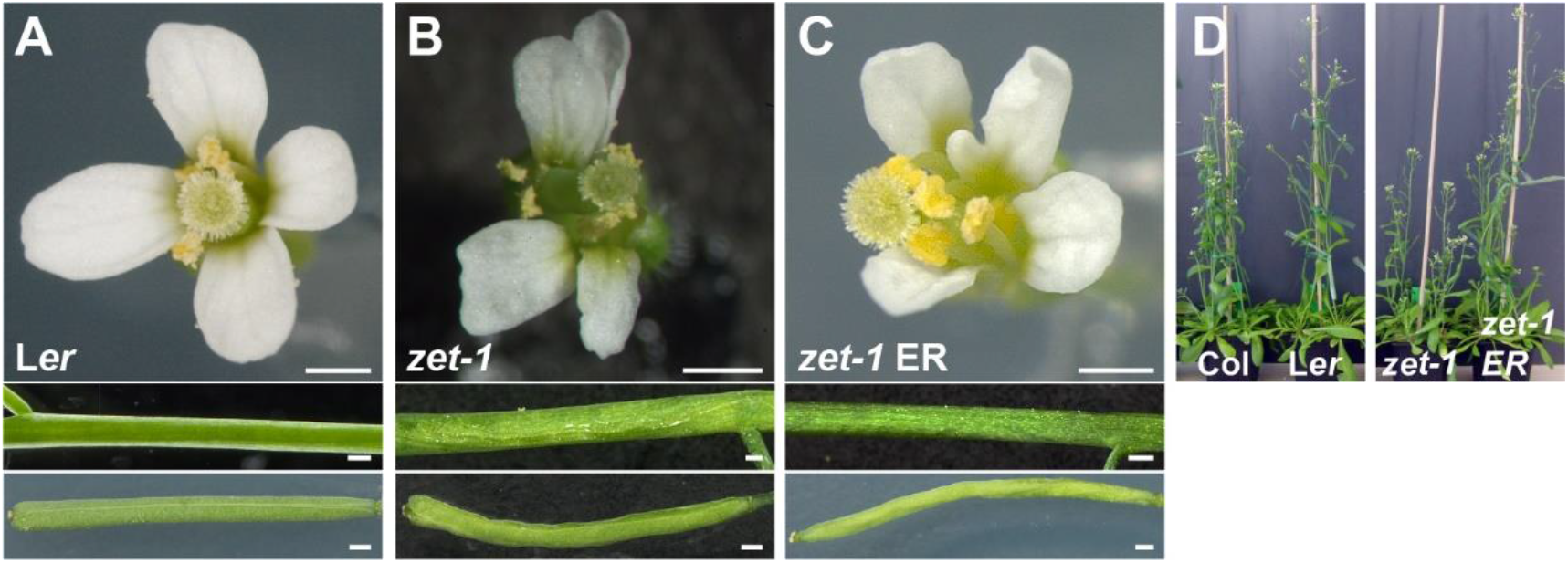
The *zet-1* phenotype in the presence of functional *ERECTA* (*ER*). (A-C) Comparison of wild type, *zet-1* and *zet-1* plants transgenic for Col *ERECTA* Morphology of flowers (upper panel), stems (central panel) and siliques (bottom panel). (A) Wild-type L*er*. (B) *zet-1*. Note the aberrant flower and silique morphology. Stem twisting is mild. (C) Transgenic *zet-1 ER*. Note the irregular flower and silique morphology. Stem morphology is essentially normal. (D) Plant height comparisons of six-week-old *zet-1 ER* transgenic plants in comparison to wild type and mutant reference lines. Note the rescue of plant height in *zet-1 ER* plants. Scale bars: (A-C) 0.5 mm

## Discussion

Numerous studies have shown the compensation of gene loss by duplicate genes implying that close homologs give robustness to the plants against mutations (Hanada *et al*. 2009). Studies have also shown stronger reduction in duplicate expression and this expression divergence was noted as an important innovation for conservation of the duplicate gene (Ganko *et al*. 2007; Panchy *et al*. 2016). Despite acknowledging this interesting pattern, examples are missing that show functional relevance of gene duplicates, since retention of almost identical duplicates goes against the evolutionary instability of genetic redundancy (Lynch and Conery 2000). Here, our work reveals an unexpected variation of a weakly expressed gene and its homolog between accessions. Our results imply that divergence in duplicate expression may play a crucial role in accession-specific genetic adaptations.

Redundancy between duplicate genes by a compensation mechanism via feedback responsive circuit serves as an advantage for biological systems to overcome stochastic fluctuations in signaling pathways (Kafri *et al*. 2006, 2009). In the Col background, an overall higher expression level of *ZETH*, in combination with a compensatory upregulation in *zet-4* mutants seems to account for their wild-type appearance. Although this phenomenon is absent in L*er*, the residual expression of *ZETH* in L*er* seems to be above the threshold, otherwise the *zet-1 zeth-2* double mutant would have resembled the single mutant *zet-1*. Furthermore, understated changes in expression pattern may further enhanced by tissue level sampling. Although segregating mutants in Col background appear to be fine morphologically, we may have missed subtler cellular phenotypes in our analysis. All the experiments were performed in controlled lab conditions. There is also a possibility that *ZETH* may express at higher levels under different conditions and reveal a novel function. Indeed, the role of SUB in coordinating cell proliferation and differentiation during leaf development was revealed only at high ambient temperature of 30 °C (Lin *et al*. 2012).

Initially, we assumed *ZETH* in L*er* background as a pseudogene, since a previous RNA-seq analysis has found that *ZETH* transcripts were undetectable in young L*er* flowers (Jiao and Meyerowitz 2010). Although our qRT-PCR results showed *ZETH* expression, the functionality was still in question given the very low expression pattern in all the tissues tested. But, surprisingly our results indicate that weakly expressed *ZETH* is functionally very relevant. Our results also highlight the importance of gene specific analysis, since in large scale studies differentiating between identical duplicate sequences is challenging. Thus, small expression differences are often overlooked as the focus goes inadvertently on highly expressed genes.

Previous evidence indicated that *slm*s may have additional functions apart from their role in plant development (Fulton *et al*. 2009). Transcriptome analysis showed that several *ZET* responsive genes are related to biotic and abiotic stress responses. It is intriguing to speculate that the high expression of *ZET* might be attributed to its stress related functions since minimal *ZETH* expression in Col was enough for wild type appearance of *zet-4* mutants.

Future experiments could test this possibility by assessing if *zet* is more susceptible to stress compared to *zeth*. Interestingly, *ZET* transcript level was shown to be altered in Arabidopsis plants infected with *Fusarium oxysporum* (Fallath *et al*. 2017). Thus, further analysis of *ZET* may reveal its potential role in adaptation, apart from morphogenesis.

Duplicate genes provide mutational robustness to living organisms. In yeast and *Caenorhabditis elegans*, functional compensation by the duplicated gene displayed higher robustness to gene perturbation than singletons (Gu *et al*. 2003; Conant and Wagner 2004). But, *ZETH* showed a highly reduced expression pattern compared to *ZET*. Such a reduction in expression was proposed to facilitate the retention of duplicates and the conservation of their ancestral functions (Qian *et al*. 2010). In this scenario, loss of either of the duplicate genes renders the total expression level lower than normal which would hamper the function. This also inhibits functional divergence of duplicated genes and helps in rebalancing gene dosage after duplication. Interestingly, with *ZET* this phenomenon was observed only in L*er*, where mutating *ZET* was enough for the manifestation of phenotype, but not in the Col background. These results indicate that the divergent behavior of duplicates may vary depending on the accession and the specific gene pair under study. Since the *ER* locus was able to partially alleviate *zet-1* mutant phenotype, the influence of accession-specific differences on gene contribution needs to be considered. Further, it would be interesting to know if there exists a correlation between accession-specific modifiers and expression divergence of certain duplicates.

A study on homologs revealed significant diversity in expression pattern among different accessions of Arabidopsis (Kliebenstein 2008). For instance, background specific regulation and unequal genetic redundancy has been observed for *BRI1* (Caño-Delgado *et al*. 2004; Zhou *et al*. 2004). In another example, the sucrose transporter AtSUC1 was shown to have differential tissue expression pattern depending on the accession it was tested (Feuerstein *et al*. 2010). The observed accession-specific disparities in *ZETH* expression levels may relate to differences in its *cis*-regulatory region, either caused by DNA polymorphisms, as was found for the tomato *fw2.2*, rice *qSH1*, or Arabidopsis *FLOWERING LOCUS* (*FT*) loci (Konishi *et al*. 2006; Cong *et al*. 2008; Schwartz *et al*. 2009; Liu *et al*. 2014), or by epigenetic variation (Durand *et al*. 2012). Thus, our work provides an interesting example for the diversification of cis and/or trans regulatory elements between two Arabidopsis accessions. A correlation between duplicate divergence and evolution of cis-regulatory elements and networks was observed (Arsovski *et al*. 2015). But, how the transcriptional networks regulate the levels of the duplicated genes and their role in the context of evolution is largely unexplored. Further studies, using the various Arabidopsis accessions available, could help in understanding the role of expression divergence among duplicates in differentiation of accessions.

## Materials and Methods

### Plant work, Plant Genetics and Plant Transformation

*Arabidopsis thaliana* (L.) Heynh. var. Columbia (Col-0) and var. Landsberg (*erecta* mutant) (L*er*) were used as wild-type strains. Plants were grown as described earlier (Fulton *et al*. 2009). The *zet-1* mutant was described previously (Vaddepalli *et al*. 2017). T-DNA insertion lines were received from the GABI-KAT (*zet-3*, GABI-KAT-460G06) (Kleinboelting *et al*. 2012) and Wisconsin collections (*zet-4*, WiscDsLoxHs057_03H; *zeth-1*, WiscDsLoxHs066_12G) (Sussman *et al*. 2000). Plants were transformed with different constructs using Agrobacterium strain GV3101/pMP90 (Koncz and Schell 1986) and the floral dip method (Clough and Bent 1998). Transgenic T1 plants were selected on Hygromycin (20 μg/ml) or Glufosinate (Basta) (10 μg/ml) plates and transferred to soil for further inspection.

### Recombinant DNA work

For DNA and RNA work standard molecular biology techniques were used. PCR-fragments used for cloning were obtained using Phusion high-fidelity DNA polymerase (New England Biolabs, Frankfurt, Germany) or TaKaRa PrimeSTAR HS DNA polymerase (Lonza, Basel, Switzerland). PCR fragments were subcloned into pLitmus 28i (NEB). All PCR-based constructs were sequenced. Primer sequences used for cloning and qRT PCR in this work are listed in S1 Table.

### Cloning

Genomic fragments of *ZETH* were amplified from Col-0 and *Ler* backgrounds using primers ZETHCol_F/ZETHCol_R and ZETHL*er*_F/ZETHL*er*_R and sub cloned into pZET::TS:ZET (Vaddepalli *et al*. 2017) using XmaI/BamHI restriction sites to obtain pZET::TS:ZETHCol and pZET::TS:ZETHL*er* respectively. For CRISPR/cas9 ZETH construct, the egg cell-specific promoter-controlled CRISPR/Cas9 system was used as described (Wang *et al*. 2015). *zet-1* plants were transformed with the CRISPR/cas9 ZETH construct by floral dip method. T1 plants were screened for exaggerated phenotype and *ZETH locus* was sequenced for identifying the mutation.

### Quantitative RT-PCR analysis

Tissue for quantitative real-time PCR (qPCR) was harvested from plants grown in long day conditions. With minor changes, tissue collection, RNA extraction and quality control were performed as described previously (Box *et al*. 2011). RT-PCR was performed on Biorad CFX96 by using iQ SYBR Green Supermix (Bio-Rad) according to the manufacturer's recommendations. All expression data were normalized against reference genes At5g25760, At4g33380, and At2g28390 by using the ΔΔ-Ct method (Czechowski 2007). Experiments were performed in biological and technical triplicates.

## Acknowledgements

We acknowledge Maxi Oelschner for technical assistance. We also thank Dolf Weijers and Sumanth Mutte for their constructive feedback on the manuscript. This work was funded by the German Research Council (DFG) through grants SCHN 723/6-1 and SFB924 (TP A2) to KS.

## Author Contributions

P.V. and K.S. designed the research. P.V and L.F. performed experiments. P.V. and K.S. analyzed the data. P.V. wrote the paper.

---

**Fig. S1.**
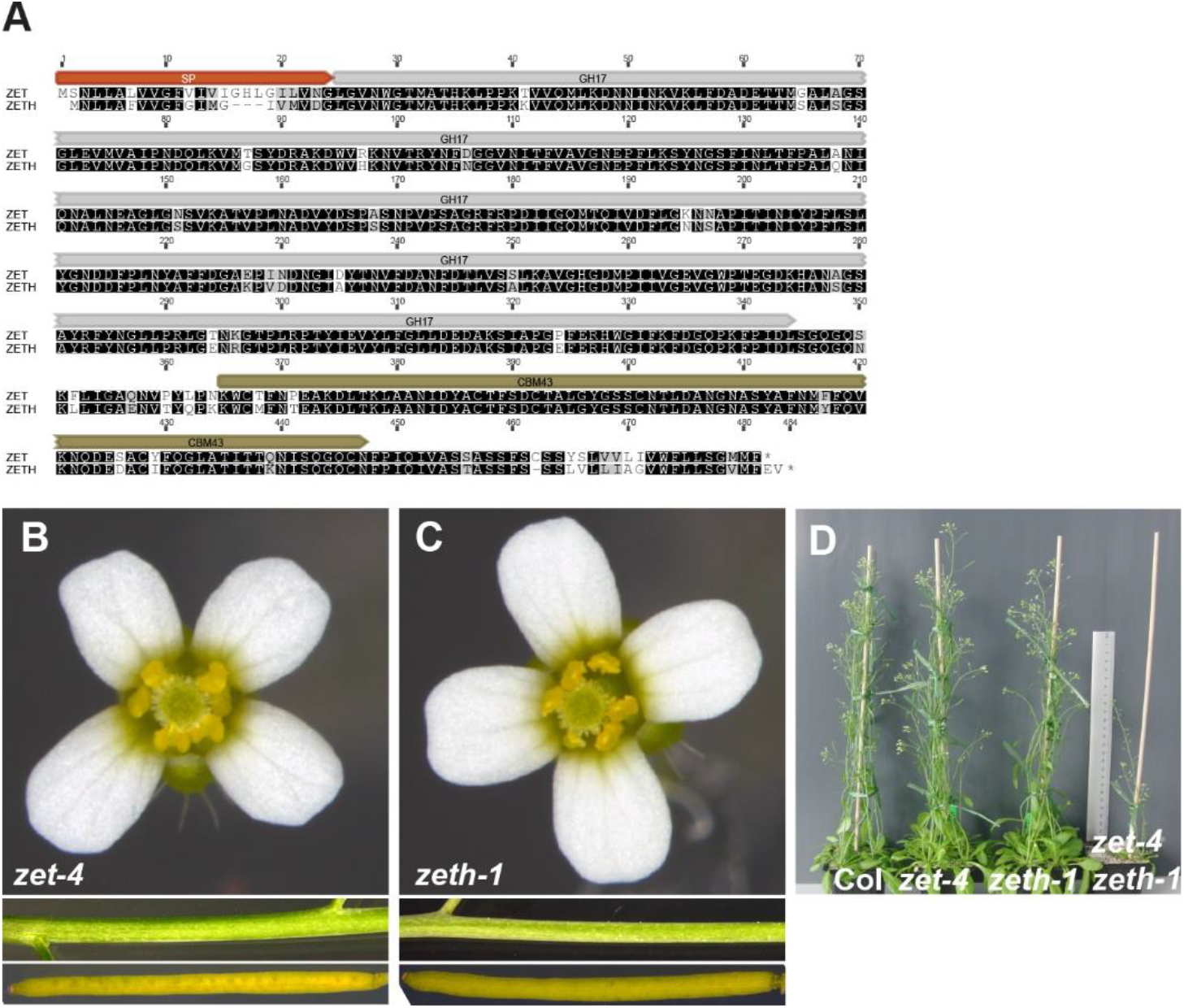
A) Alignment of ZET and ZETH amino acid sequences. B) *zet-4*. C) *zeth-1*. Flower, stem and silique morphology is essentially normal. (D) Whole-plant appearance. Genotypes are indicated.

**Fig. S2.**
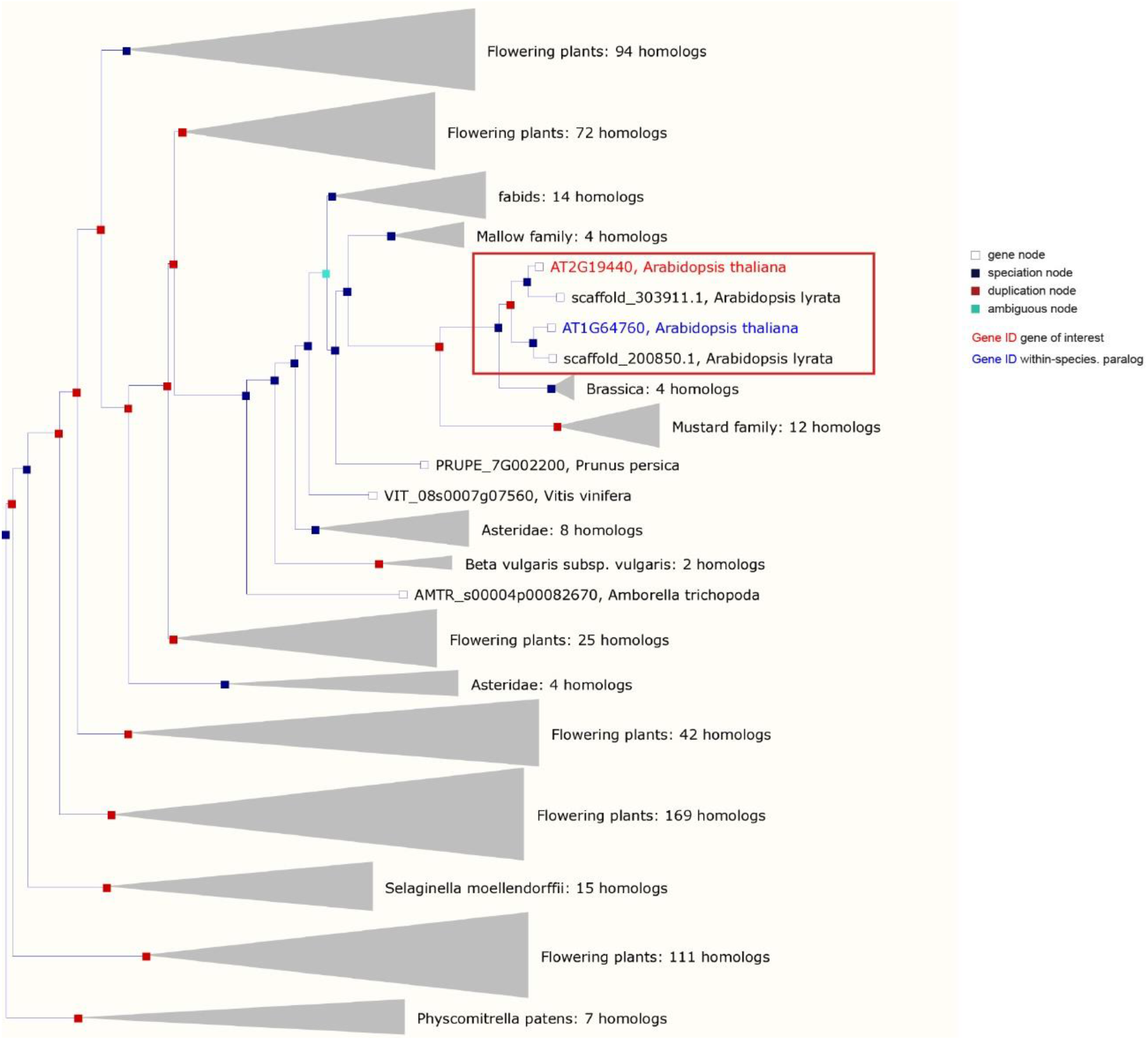
Evolutionary history of ZET (AT1G64760) and ZETH (AT2G19440) Phylogenetic tree showing the evolution of the ZET and its closest homolog (ZETH). The tree shows that duplication is specific to Arabidopsis and it is not present in any other Brassicaceae members. (Source: http://plants.ensembl.org)

**Supplementary Table S1.**
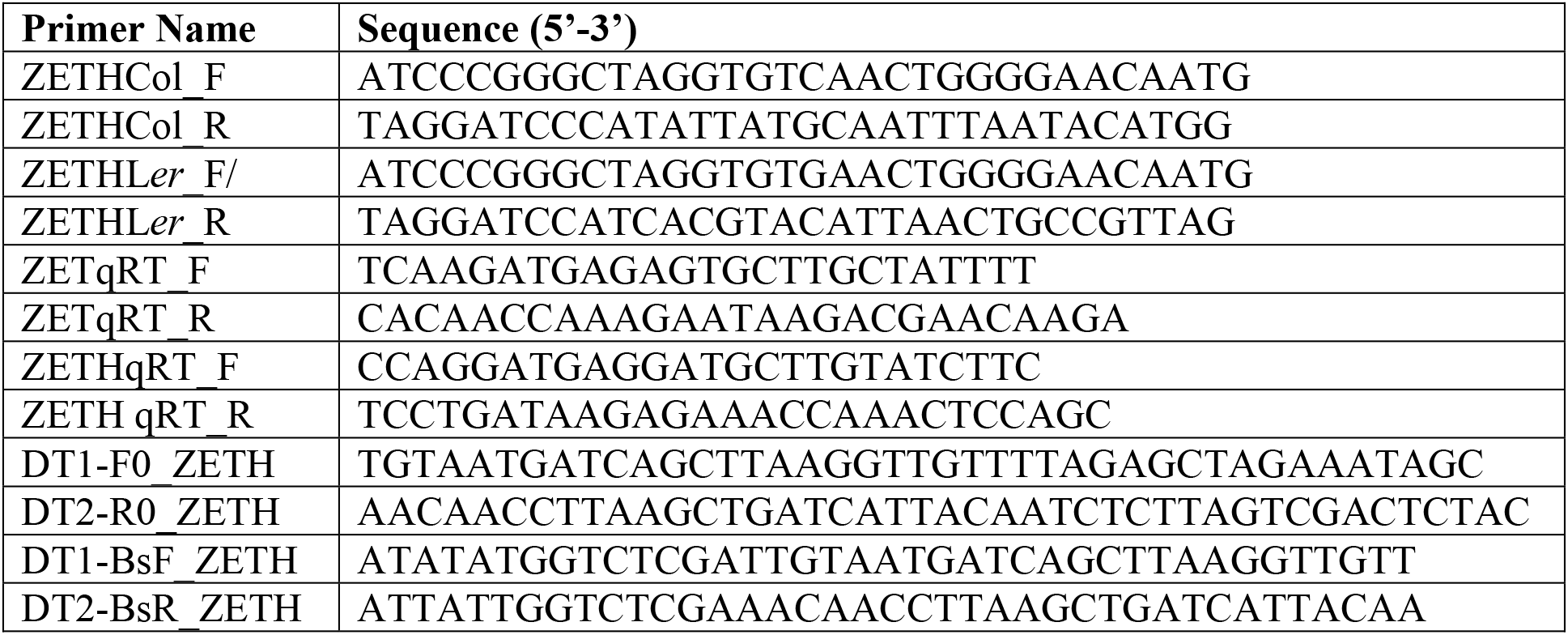
Primers used in this study

